# Mind Wandering in Sensory Cortices

**DOI:** 10.1101/2020.04.27.064360

**Authors:** Shao-Min Hung, Po-Jang Hsieh

## Abstract

Mind wandering contains rich phenomenology as we experience moment by moment, however, such linkage between our subjective experiences and the underlying neural mechanism has been missing in the literature. Here we report that the sensory contents of mind wandering recruit corresponding sensory cortices, serving as the neural bases of the sensory contents in mind wandering.

## Introduction

Mind wandering occupies a significant amount of our waking time(*1*) and interferes with our task performance(*2*). Despite its prevalence, the functional significance of mind wandering is still under debate(*3*). Recent research has begun to reveal the neural mechanisms underlying mind wandering, specifically pointing to a linkage between mind wandering and the default mode network(*4, 5*) (DMN) and other networks such as the executive system(*5*). The DMN, including the medial prefrontal cortex, tempero-parietal junction, and posterior cingulate cortex, is most active when humans are at rest or off-task(*6*), shedding light on the *functional* nature of mind wandering as a default system which our mind regresses to. However, the *phenomenal* nature of mind wandering has been largely missing in the neuroimaging literature, leaving a huge unexplanatory gap between imaging findings and our rich sensory experiences during mind wandering. As mind wandering accompanies rich phenomenology, we expected to observe corresponding sensory experiences to recruit sensory cortices.

## Results

Here we introduced online thought sampling(*7*) in an MRI study to capture and categorize the sensory contents during mind wandering. Three participants each underwent 10 hours of scanning in 5 separate sessions during which the task was to fixate on a dot. In a run lasting 8 minutes, the participants were probed at random intervals (45 ∼ 90 s), with the intervals optimized based on the frequency of mind wandering individually. The first probe asked the participants if they were on-task, which denoted that they were focused on the fixation task without having task-irrelevant thoughts. If off-task report was given, a series of questions regarding the mind wandering contents were asked to quantify the mind wandering event immediately preceding the probe (Fig. 1). In total over 200 mind wandering events were collected for each participant.

**Fig. 1.**
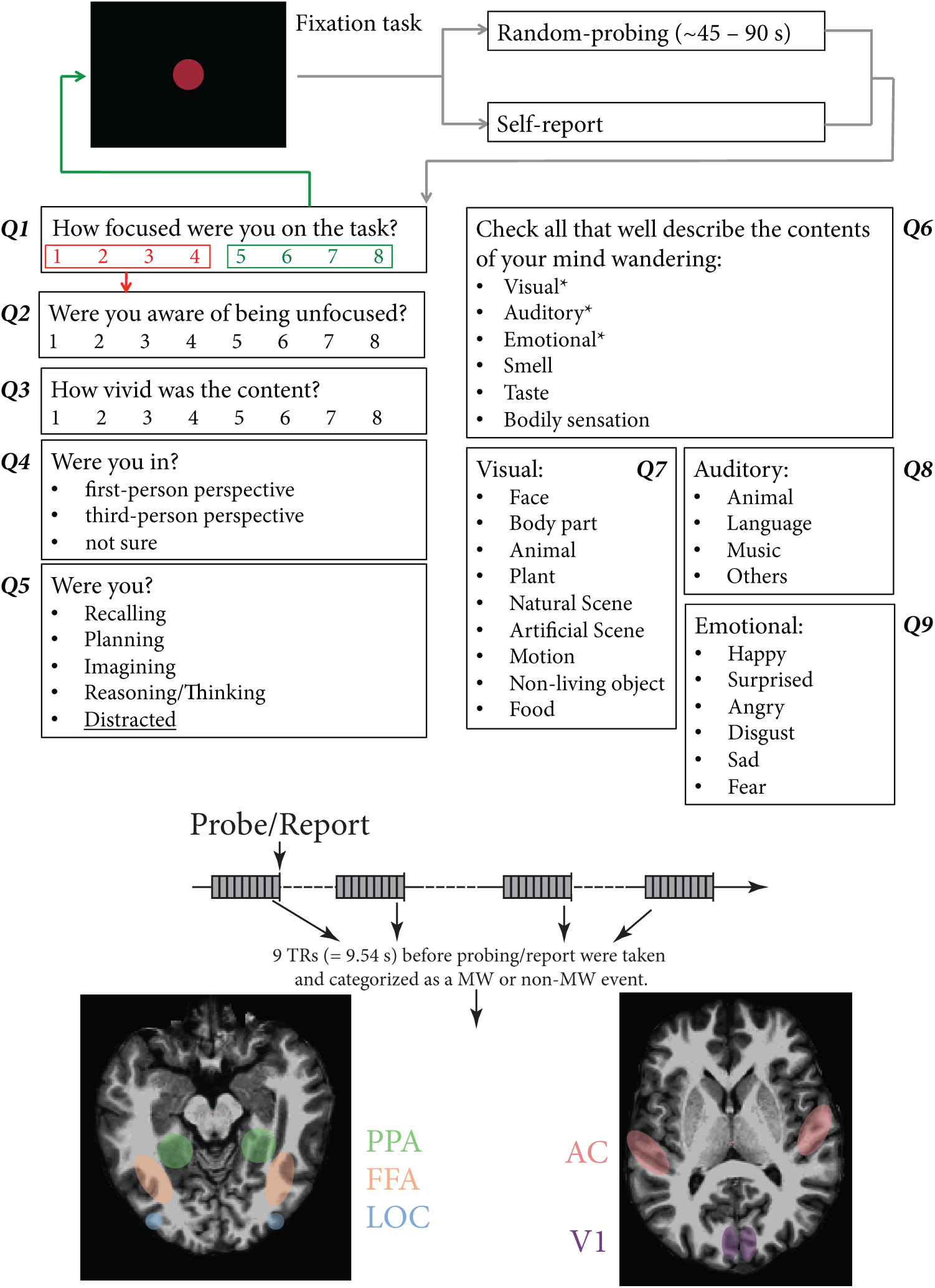
Online content sampling procedure. **Top**. The participant underwent a fixation task in each 8-min run. Every 45 ∼ 90 seconds, adjusted according to each individual’s mind wandering frequency, a question popped out and asked Q1: How focused were you on the task? If the answer was 5 ∼ 8, this probe was categorized as a non-mind wandering event, and the participant resumed the fixation task immediately. If the answer was 1 ∼ 4, the participant proceeded to report a series of questions regarding the mind wandering contents. In Q6, if either of the three answers (visual/auditory/emotional) was chosen, a corresponding subsequent subcategory question would be asked to gather the details of sensory content. Participants were allowed to report their mind wandering event if they caught it voluntarily, however, no such event took place in our study. **Bottom**. 9 TRs precedent to the probe was categorized as a non-mind wandering or a wandering event and labelled with respective sensory contents. Five regions including Auditory Cortex (AC, in pink), striate cortex (V1, in purple), FFA (in orange), LOC (in blue), and PPA (in green) were functionally localized in each individual in a separate scan. The analyses were performed in these sensory ROIs.

9 TRs (9.54 s) before the probe were taken and labeled as a mind wandering or non-mind-wandering event. The mind-wandering events were further labelled with exclusive specific sensory categories (e.g. visual/auditory) and the subcategories (e.g. face/object/scene/etc.). In a separate localizer scan, the participants underwent retinotopic mapping, audio localizer, and visual localizer to individually localize striate cortex (V1), auditory cortex (AC), the face fusiform area (FFA), the parahippocampal place area (PPA), and the lateral occipital complex (LOC). Further analyses were performed solely on these pre-determined functional ROIs. In each mind wandering event, the beta values of all the voxels within an ROI were extracted. Before submitted these event vectors to support vector machine (SVM), all the vectors were normalized against the mean beta value. Half of the event vectors were used to train a linear classifier while the other half were used for testing. All comparisons were made pairwise. One critical limitation of the current study, which will be faced by almost any future mind wandering studies, is the impulsive and uncontrollable nature of mind wandering episodes. This factor posed a major deficit on the decoding technique as one might be decoding the nuanced differences between two sensory contents, such as distinct temporal occurrence. In order to directly tease apart the decoding performance that was contributed by sensory content differences and these nuanced differences, we randomly shuffled the labels of these sensory contents and performed the same SVM analysis. The performance of these random shuffling decoding accuracy is deemed the true baseline accuracy.

Two major comparisons with enough exclusive sensory contents (e.g. visual-only) were made. In mind wandering events involved visual-only versus auditory-only contents, we showed successful decoding in these ROIs (Fig. 2). Importantly, the profile in each individual varied significantly, with some individual showing above-chance decoding accuracy in all visual and auditory ROIs while some individual showing better decoding performance only in a high-level region (i.e. LOC). In mind wandering events involved face-only versus object-only contents, we utilized PPA as the baseline region and expected to see chance performance. Again, individuals show distinct decoding patterns in FFA and LOC, underscoring the vast individual differences of subjective phenomenology in mind wandering. Across all individuals, successful decoding in FFA was observed with different participants showing distinct decoding accuracies in LOC. The more consistent decoding performance in FFA could indicate the strong bias of mind wandering content towards human faces (Table 1) and less variance in physical properties across distinct faces, as compared to objects in this pairwise comparison.

**Table 1.** Detailed individual online mind wandering reports. In total, approximately 350 probes occurred in the span of 50 8-min runs, in which over 200 mind wandering events were collected. Most reports had overlapped sensory contents, while the reports with exclusive sensory contents are shown in parentheses and used for analyses. Asterisk denotes contents with further subcategories.

|  | MW 01 | MW 02 | MW 03 |
| --- | --- | --- | --- |
| Total probes | 346 | 368 | 349 |
| MW events | 233 | 231 | 264 |
| Non-MW events | 113 | 137 | 85 |
| MW percentage | 67.3 % | 62.8 % | 75.6 % |
| Sensory categories |  |  |  |
| Visual* | 183 (24) | 139 (21) | 196 (40) |
| Auditory* | 115 (11) | 83 (17) | 108 (14) |
| Emotional* | 87 (9) | 42 (5) | 44 (2) |
| Bodily | 24 (6) | 35 (11) | 40 (8) |
| Taste | 3 (0) | 10 (1) | 17 (1) |
| Conceptual | 80 (6) | 120 (46) | 57 (33) |
| Smell | 1 (0) | 7 (0) | 18 (0) |
| Visual subcategories |  |  |  |
| Face | 132 (25) | 74 (12) | 109 (17) |
| Body | 32 (0) | 50 (4) | 102 (7) |
| Animal | 5 (1) | 0 (0) | 14 (5) |
| Plant | 6 (1) | 1 (0) | 3 (0) |
| Natural scene | 27 (3) | 5 (0) | 13 (1) |
| Artificial scene | 99 (14) | 75 (7) | 12 (0) |
| Motion | 14 (4) | 25 (2) | 21 (0) |
| Object | 34 (13) | 52 (14) | 78 (26) |
| Food | 5 (2) | 11 (1) | 29 (7) |
| Auditory subcategories |  |  |  |
| Animal | 1 | 2 | 3 |
| Language | 94 (74) | 66 (54) | 100 (94) |
| Music | 39 (19) | 19 (14) | 8 (2) |
| Others | 3 | 1 | 9 |
| Emotion subcategories |  |  |  |
| Happy | 40 (25) | 18 (14) | 15 (15) |
| Surprised | 12 (8) | 0 (0) | 14 (13) |
| Angry | 13 (8) | 5 (3) | 6 (5) |
| Disgusted | 7 (3) | 3 (2) | 5 (5) |
| Sad | 12 (8) | 0 (0) | 14 (13) |
| Fear | 23 (17) | 5 (5) | 1 (1) |

**Fig. 2.**
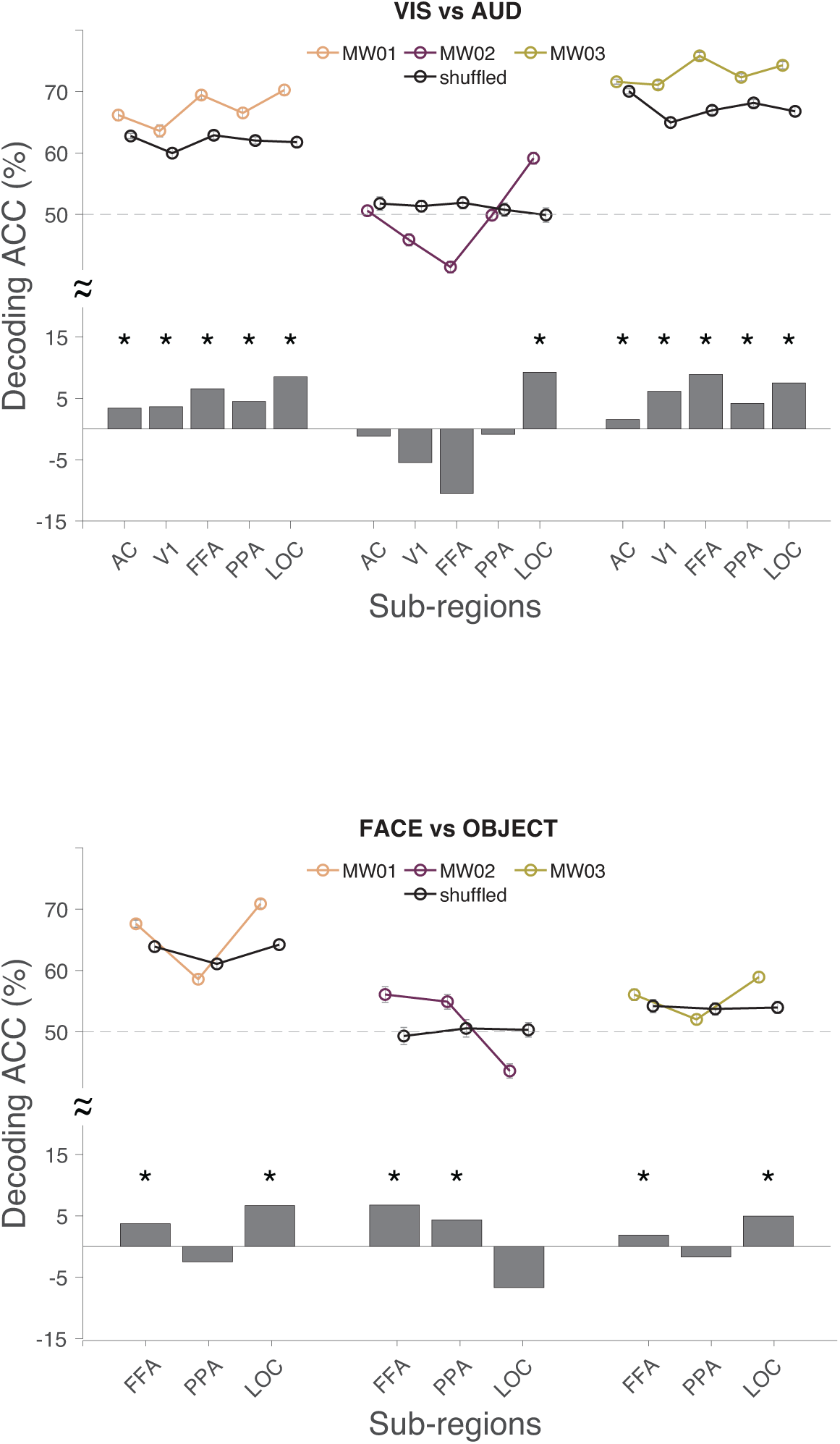
Decoding accuracy (ACC) of three individuals in sensory ROIs. The decoding ACC was compared against the baseline where the labels were randomly shuffled. The top half of the figure denotes decoding ACC while the bottom half shows the decoding ACC differences between correctly versus randomly labelled in bar plot. **Top**. Decoding ACC of visual-only vs. auditory-only sensory events in 5 respective sensory regions. **Bottom**. Decoding ACC of face-only vs. object-only sensory events in 3 respective sensory regions. Asterisk denotes significance (all paired t, *p* < 0.00001). Error bars denote 99.9 % confidence intervals.

## Discussion

Our study serves as one of the first brain imaging studies to directly probe the phenomenology of mind wandering, showing that distinct sensory cortices are indeed recruited under distinct mind wandering sensory contents. Importantly, each individual exhibits a unique profile of sensory cortices involvement. Such individual differences outline several key features of mind wandering. For example, the content itself could have varied significantly in each individual(*8*). Furthermore, although in the mind wandering events extracted in our study, no vividness difference was observed across individuals, in a post-study vividness of visual imagery questionnaire, these individuals did show differences (eyes closed, MW 01: 72; MW 02: 59; MW 03: 69, VVIQ(*9*)), suggesting the ability to generate imagery possibly varied. In fact, a previous study has shown concurrent suppression of auditory cortex activation during active visual imagery(*10*). Importantly, such suppression negatively correlated with VVIQ scores, indicating that people with vivid visual imagery showed stronger concurrent suppression on the irrelevant auditory processing. Our data suggest that similar modality-separation mechanism could occur in mind wandering as participants with higher VVIQ scores show better decoding performance differentiating visual versus auditory contents in the corresponding visual and auditory regions. A large-scale study is needed to directly assess the interplay between different modalities and sensory contents in mind wandering, and how individual propensity to generate mind wandering plays a role. Instead of seeing individual differences as a noisy factor(*11*) in a typical imaging study, we highlight the importance of such individual-based experiment and analysis, especially when the core research interests concern personal, private subject experiences. Recently, a single-subject study(*12, 13*) conducted over 1 year period has documented dynamic changes particularly in the functional connectivity of the visual and somato-motor networks in the resting-state scan, which is similar to the mind wandering condition in the current study. Critically, this finding is largely inconsistent with an inter-subject study that showed least variability in the sensory-motor and visual regions(*14*).

Current study combines online thought sampling and brain imaging, extending our neuronal understanding of mind wandering from “functional” to “phenomenological.” As the neuroscientific study of mind wandering has bloomed in the past decades, the research has advanced tremendously and revealed general brain networks underlying this involuntary subjective experience(*15*). Our study further compliments the picture of mind wandering study by showing the neuronal bases of the rich phenomenology under our daily mind wandering.

## Materials & Methods

### Participants

Three participants (age range: 25 ∼ 29) from the Duke-National University of Singapore Medical school community were recruited (1 male) and took part in the six-session study. Each session lasted 2 hours. All participants reported free of any neurological, psychiatric, and sleep disorders. They had normal or corrected-to-normal vision. The experiments were approved by the institutional review boards at the National University of Singapore. All participants gave written informed consent prior to the experiments and were reimbursed with $35/session.

### fMRI experiment

#### Design

Scanning was performed using a 3T Siemens Prisma scanner (Siemens, Erlangen, Germany) at the Duke-NUS Medical School, Singapore. Functional MRI runs were acquired using a gradient echo-planar imaging multiband sequence (TR 1.06 s, TE 32 ms, FA 61°, FOV 1980 × 1980 mm, 2 × 2 mm in-plane resolution). Thirty-six slices were collected with a 12-channel head coil (2.0 mm thickness). Slices were oriented roughly parallel to the AC-PC with whole brain covered. A T1-weighted anatomical image was also acquired and later used for co-registration (TR 2.3 s, TI 900 ms, FA 8°, FOV 256 × 240 mm, 192 slices, 1×1×1 mm). Each participant took part in approximately 10 runs per session with 5 sessions in total for the main mind wandering experiment. An additional functional localizer session was collected. To localize the striate cortex (V1), three runs of retinotopic mapping scan were run. Retinotopic mapping consisted of six 20-s blocks each flanked by 20-s fixation. Stimuli were presented in three experimental conditions. Each condition was repeated twice in a single run. The conditions were presented in a pseudo-randomized order across all three blocks. In the retinotopic mapping scans, flashing checkerboard wedges were presented in each condition. In the horizontal condition, two wedges subtending 10° from the central fixation were presented along the horizontal meridian. Similarly, in the vertical condition, the two wedges were presented along the vertical meridian. In the last experimental condition, four wedges each subtending 30° from fixation were presented along the diagonal axis. During a stimulus block, color of the central fixation changed between green and red. Subjects were tasked to maintain fixation at all times and indicate color of central fixation cross via button presses(*16*).

Functional localization of three of the regions of interest (ROIs) was based on four independent runs of 20-s blocks with grayscale images of faces, scenes, common objects and scrambled objects (four blocks per category per run, followed(*17*)). The fusiform face area (FFA(*18*)) was defined as the region of the fusiform gyrus that responded more strongly to images of faces than to images of intact scenes. The parahippocampal place area (PPA(*19*)) was defined as the region of the parahippocampal gyrus that responded more strongly to images of scenes than to images of intact faces. Similarly, the lateral occipital complex (LOC(*20*)) was defined as the region responded more strongly to images of intact objects than to those of scenes. All statistical maps were corrected with cluster-thresholding (*p* < 0.05; cluster–forming threshold *p* < 0.01).

Participants were instructed to do an 8-min fixation task in each experiment run while not thinking about anything in particular. They were told that anything irrelevant to the fixation task will be regarded as mind wandering. Two probed were implemented: Random probing occurred every 45 ∼ 90 seconds while participants could report mind wandering anytime if they became aware of it. Please note that although we allowed self-report in the current paradigm, due to our high sampling rate, all participants never became aware of the mind wandering before the probe. The random probing interval was adjusted according to the run-by-run performance with the goal to catch 1 mind wandering event per minute. In total 350 probe occurred over 50 runs in all participants with approximately 70% of them mind wandering events. Objectively, whether participants mind wandered depended on an 8-point scale of the first probe question (“How focused were you on the task?”): 1 – 4 were deemed a mind wandering event leading to a series of questions documenting the content of mind wandering, while 5 – 8 were labelled as a non-mind-wandering event and the participant returned to the fixation task immediately (Fig. 1). A detailed table of individual reports are shown in Table 1.

### Data preprocessing

fMRI data analysis was conducted using *freesurfer* (http://surfer.nmr.mgh.harvard.edu/) and MATLAB (The MathWorks, Inc., Natick, MA). The processing steps for both the retinotopic mapping and the experimental runs included motion correction and linear trend removal. The processing for the retinotopic mapping and localizer runs used for later univariate (mean) BOLD responses analysis also included spatial smoothing with a 6-mm kernel. For every participant, all the localizer runs were modeled with general linear model (GLM). A gamma function with delta (δ)=2.25 and tau (τ)=1.25 was used to estimate the hemodynamic response for each condition in the retinotopic mapping and localizer scans and the experimental scans. For the experimental runs, the time courses were obtained with a finite impulse response (FIR) model without assuming a particular hemodynamic response function. Such FIR model has been used to identify pre-trial signals(*21*).

### Mind wandering event labeling and multivariate pattern analysis

The β value of 9 TRs (9.54 s) preceding probe or report were extracted and labeled as a mind wandering or a non-mind-wandering event (Fig. 2, procedure similar to(*7*)). Each mind wandering event was further labeled with corresponding content according participants’ online report. We chose two major comparisons: visual vs. auditory and face vs. object vs. scenes. Please note that these events were exclusive to one category. For instance, visual events were the events that participants reported only visual contents with no other sensory contents (e.g. face only). Multivariate pattern analysis was performed in each corresponding ROI of each participant. The β value in each voxel was extracted in each comparing condition in the designated ROI. Furthermore, to ensure mean activation difference would not contribute to our multivariate analysis result, β value in each voxel in each condition was normalized against the mean response of the condition before further analysis. A binary linear support vector machine (MATLAB, *fitcsvm*) was built for each comparison. Half of the events from the two comparing conditions were used to train the linear classifier, while the other half were reserved for later testing. Each time an accuracy rate was obtained by dividing the hit votes with total testing votes. This iteration was repeated 1000 times for each comparison. Later the mean decoding accuracy and 99.9% confidence interval were derived across all iterations.

## AUTHOR CONTRIBUTIONS

Conceptualization, S.-M.H. and P.-J. H.; Methodology, S.-M.H. and P.-J. H.; Software, S.-M.H.; Investigation, S.-M.H.; Writing – Original Draft, S.-M.H.; Writing – Review & Editing, S.-M.H. and P.-J. H.; Supervision, P.-J. H.

## ACKNOWLEDGEMENTS

This work was supported by the Yushan Young Scholar Program (NTU-108V0202). We thank the support of James Boswell Postdoctoral Fellowship and Caltech Divisional Postdoctoral Fellowship to S.-M.H.

## References

1. M. A. Killingsworth, D. T. Gilbert, A Wandering Mind Is an Unhappy Mind. Science. 330, 932–932 (2010).

2. J. C. McVay, M. J. Kane, Conducting the train of thought: Working memory capacity, goal neglect, and mind wandering in an executive-control task. Journal of Experimental Psychology: Learning, Memory, and Cognition. 35, 196–204 (2009).

3. J. Smallwood, J. W. Schooler, The Science of Mind Wandering: Empirically Navigating the Stream of Consciousness. Annu. Rev. Psychol. 66, 487–518 (2015).

4. M. F. Mason, M. I. Norton, J. D. Van Horn, D. M. Wegner, S. T. Grafton, C. N. Macrae, Wandering Minds: The Default Network and Stimulus-Independent Thought. Science. 315, 393–395 (2007).

5. K. Christoff, A. M. Gordon, J. Smallwood, R. Smith, J. W. Schooler, Experience sampling during fMRI reveals default network and executive system contributions to mind wandering. Proceedings of the National Academy of Sciences. 106, 8719–8724 (2009).

6. R. L. Buckner, J. R. Andrews-Hanna, D. L. Schacter, The Brain’s Default Network: Anatomy, Function, and Relevance to Disease. Annals of the New York Academy of Sciences. 1124, 1–38 (2008).

7. T. Horikawa, M. Tamaki, Y. Miyawaki, Y. Kamitani, Neural Decoding of Visual Imagery During Sleep. Science. 340, 639–642 (2013).

8. D. Marcusson-Clavertz, E. Cardeña, D. B. Terhune, Daydreaming style moderates the relation between working memory and mind wandering: Integrating two hypotheses. Journal of Experimental Psychology: Learning, Memory, and Cognition. 42, 451–464 (2016).

9. D. F. Marks, VISUAL IMAGERY DIFFERENCES IN THE RECALL OF PICTURES. British Journal of Psychology. 64, 17–24 (1973).

10. A. Amedi, R. Malach, A. Pascual-Leone, Negative BOLD Differentiates Visual Imagery and Perception. Neuron. 48, 859–872 (2005).

11. R. Kanai, G. Rees, The structural basis of inter-individual differences in human behaviour and cognition. Nat Rev Neurosci. 12, 231–242 (2011).

12. R. A. Poldrack, T. O. Laumann, O. Koyejo, B. Gregory, A. Hover, M.-Y. Chen, K. J. Gorgolewski, J. Luci, S. J. Joo, R. L. Boyd, S. Hunicke-Smith, Z. B. Simpson, T. Caven, V. Sochat, J. M. Shine, E. Gordon, A. Z. Snyder, B. Adeyemo, S. E. Petersen, D. C. Glahn, D. Reese Mckay, J. E. Curran, H. H. H. Göring, M. A. Carless, J. Blangero, R. Dougherty, A. Leemans, D. A. Handwerker, L. Frick, E. M. Marcotte, J. A. Mumford, Long-term neural and physiological phenotyping of a single human. Nat Commun. 6, 8885 (2015).

13. T. O. Laumann, E. M. Gordon, B. Adeyemo, A. Z. Snyder, S. J. Joo, M.-Y. Chen, A. W. Gilmore, K. B. McDermott, S. M. Nelson, N. U. F. Dosenbach, B. L. Schlaggar, J. A. Mumford, R. A. Poldrack, S. E. Petersen, Functional System and Areal Organization of a Highly Sampled Individual Human Brain. Neuron. 87, 657–670 (2015).

14. S. Mueller, D. Wang, M. D. Fox, B. T. T. Yeo, J. Sepulcre, M. R. Sabuncu, R. Shafee, J. Lu, H. Liu, Individual Variability in Functional Connectivity Architecture of the Human Brain. Neuron. 77, 586–595 (2013).

15. K. Christoff, Z. C. Irving, K. C. R. Fox, R. N. Spreng, J. R. Andrews-Hanna, Mindwandering as spontaneous thought: a dynamic framework. Nat Rev Neurosci. 17, 718–731 (2016).

16. S.-M. Hung, D. Milea, A. V. Rukmini, R. P. Najjar, J. H. Tan, F. Viénot, M. Dubail, S. L. C. Tow, T. Aung, J. J. Gooley, P.-J. Hsieh, Cerebral neural correlates of differential melanopic photic stimulation in humans. NeuroImage. 146, 763–769 (2017).

17. Z. Kourtzi, N. Kanwisher, Representation of Perceived Object Shape by the Human Lateral Occipital Complex. Science, New Series. 293, 1506–1509 (2001).

18. N. Kanwisher, J. McDermott, M. M. Chun, The Fusiform Face Area: A Module in Human Extrastriate Cortex Specialized for Face Perception. Journal of Neuroscience. 17, 4302–4311 (1997).

19. R. Epstein, N. Kanwisher, A cortical representation of the local visual environment. Nature. 392, 598–601 (1998).

20. R. Malach, J. B. Reppas, R. R. Benson, K. K. Kwong, H. Jiang, W. A. Kennedy, P. J. Ledden, T. J. Brady, B. R. Rosen, R. B. Tootell, Object-related activity revealed by functional magnetic resonance imaging in human occipital cortex. Proceedings of the National Academy of Sciences. 92, 8135–8139 (1995).

21. P.-J. Hsieh, J. T. Colas, N. G. Kanwisher, Pre-stimulus pattern of activity in the fusiform face area predicts face percepts during binocular rivalry. Neuropsychologia. 50, 522–529 (2012).

